# A Recombinant Multivalent Vaccine (rCpa1) Induces Protection for C57BL/6 and HLA Transgenic Mice Against Pulmonary Infection with Both Species of Coccidioides

**DOI:** 10.1101/2021.10.20.465232

**Authors:** Althea Campuzano, Komali Devi Pentakota, Yu-Rou Liao, Hao Zhang, Nathan Wiederhold, Gary Ostroff, Chiung-Yu Hung

**Affiliations:** Department of Biology, The University of Texas at San Antonio, San Antonio, TX, USA; Department of Pathology, Graduate School of Biomedical Sciences, UT Health, San Antonio, TX, USA; Program in Molecular Medicine and Department of Medicine, University of Massachusetts Medical School, Worcester, MA, USA

**Keywords:** Coccidioidomycosis, vaccine, cross-protection, T-cell immunity, Valley fever, HLA-DR4 mice

## Abstract

Coccidioidomycosis is caused by *Coccidioides posadasii* (*Cp*) and *Coccidioides immitis* (*Ci*) that have 4-5% differences in their genomic sequences. There is an urgent need to develop a human vaccine against both species. A previously created recombinant antigen (rCpa1) that contains multiple peptides derived from *Cp* isolate C735 is protective against the autologous isolate. The focus of this study is to evaluate cross-protective efficacy and immune correlates by the rCpa1- based vaccine against both species of *Coccidioides*. DNA sequence analyses of the homologous genes for the rCpa1 antigen were conducted for 39 and 17 clinical isolates of *Cp* and *Ci*, respectively. Protective efficacy and vaccine-induced immunity were evaluated for both C57BL/6 and human HLA-DR4 transgenic mice against 5 highly virulent isolates of *Cp* and *Ci*. There are a total of 7 amino acid substitutions in the rCpa1 antigen between *Cp* and *Ci*. Both C57BL/6 and HLA-DR4 mice that were vaccinated with a rCpa1 vaccine resulted in significant reduction of fungal burden and increased numbers of IFN-γ- and IL-17-producing CD4^+^ T cells in the first 2 weeks post-challenge. These data support that rCpa1 has cross-protection activity against *Cp* and *Ci* pulmonary infection through activation of early Th1 and Th17 responses.

## Introduction

Coccidioidomycosis (also known as Valley fever) is caused by two phylogenetically related fungal pathogens, *Coccidioides immitis* (*Ci*) and *C. posadasii* (*Cp*) that are endemic to the desert southwest of the United States and aerial regions of Central and South America (1). *Coccidioides* species are dimorphic fungi, growing as molds in the soil and differentiating into multinucleated spherules (∼100 μm in diameter) in mammals (2). Coccidioidomycosis typically begins as a pulmonary infection via inhalation of airborne arthrospores produced by the soil-dwelling mycelia. Arthrospores then undergo isotropic growth converting into spherules that form small compartments via septation process. Each compartment contains 2-4 nuclei and subsequently develops into endospores that are then released during the endosporulation process (∼300-800 endospores per spherule). Coccidioidal endospores (2-7 μm in diameter) can be extrapulmonary disseminated to the skin, bones, the central nerve system, as well as other organs via blood and lymphatic system. Therefore, chronic pulmonary and disseminated coccidioidomycosis patients may require life-long antifungal therapies (3).

Clinical and laboratory experimental results revealed that these two fungi are similar in virulence and dimorphic lifestyle (4). While population genomics analysis demonstrate that hybridization and genetic introgression have occurred between these species (5). The introgression direction is primarily from *C. posadasii* to *C. immitis*, respectively (5). There are diverse clinical isolates with hybrid genomic sequences (5-7). Approximately, the annotated orthologous genes shared between these two species present over 90% protein sequence identity (1, 8). However, *Cp* and *Ci* are indistinguishable via serological tests using coccidioidal complement fixation (CF) or tube precipitin (TP) antigens (4, 9, 10).

Knowledge gaps in the diagnosis of coccidioidomycosis, its treatment, immunological responses, its morbidity, the economic healthcare burden, among increased incidence have raised great concerns (11). Vaccination against coccidioidomycosis seems feasible since a secondary infection is extraordinarily rare (12). Our effort focuses on the development of a subunit vaccine composed of coccidioidal antigens plus an adjuvant delivery system (1, 13, 14). A successful vaccine candidate should confer protection against both *Coccidioides* species. This approach requires identifying conserved antigens among clinical isolates of these two fungal species that stimulate long-lasting, antigen-specific adaptive immune responses. Several studies have demonstrated that a multivalent vaccine can induce large amounts of specific T-cell clones, becoming more effective compared to an individual antigen (15-17). A <u>r</u>ecombinant <u>c</u>himeric <u>p</u>olypeptide <u>a</u>ntigen (rCpa1) was genetically engineered to link together the most immunogenic fragment of proline-rich coccidioidal antigen (Ag2/Pra), the full lengths of *Coccidioides* specific antigen (Cs-Ag) as well as proximal matrix protein 1 (Pmp1), as well as five promiscuous T-cell epitopes that have high affinity to human MHC class II molecules in a single polypeptide construct (**Fig. 1A**) (18, 19). The 5 T-cell epitopes are derived from *C. posadasii-*specific aspartyl protease (Pep1), a-mannosidase (Amn1), and phospholipase B (Plb) antigens (15, 16, 18, 20, 21). The multivalent rCpa1 antigen is then loaded into an adjuvant made of glucan-chitin particles to form the GCP-rCpa1 vaccine (18, 19). The vaccine has previously been reported to elicit a mixed CD4 T-cell mediated type I (Th1) and Th17 immunity and confers protection against autologous isolate (C735) of *C. posadasii* in murine models compared to the mock- immunized group (GCP-MSA) (18, 19). Previous studies only evaluated the protective efficacy of GCP-rCpa1 against *C. posadasii* isolate C735. However, we did not test the efficacy of our multivalent subunit vaccine against other *C. posadasii* isolates or *C. immitis*. In this study, we evaluated protective efficacies and CD4 T-cell mediated immunity of the GCP-rCpa1 vaccine against pulmonary infection with various isolates of *C. posadasii* and *C. immitis* using C57BL/6 and human HLA-DR4 transgenic mouse model of coccidioidomycosis.

**Figure 1.**
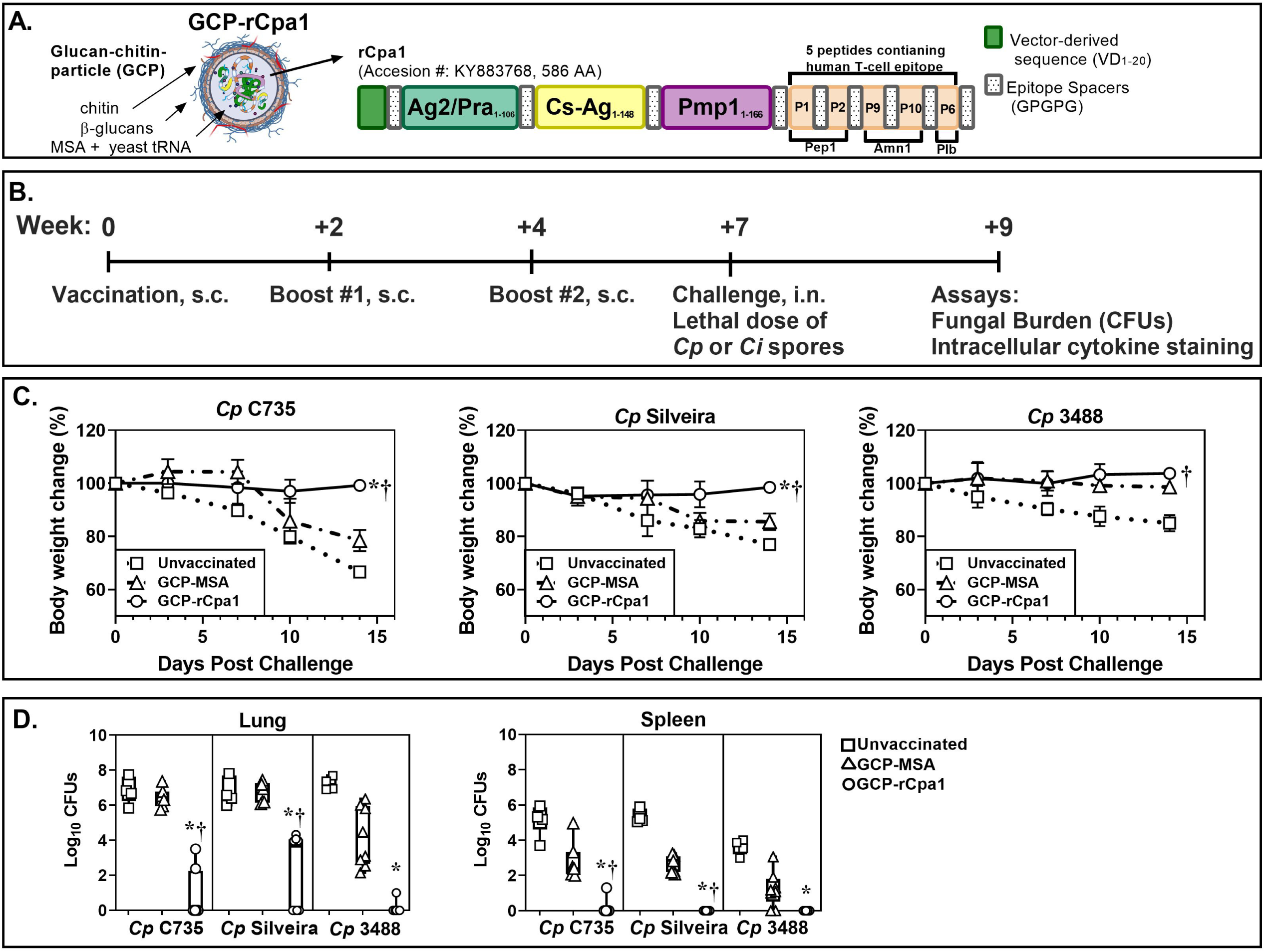
GCP-rCpa1 vaccination results in protection against multiple clinical isolates of *Cp*. (A) GCP-rCpa1 vaccine construct illustration. (B) Timeline shows vaccination and immunological analysis schedule. (C) The daily body weight change (%) of C57BL/6 mice that were subcutaneously vaccinated with the GCP-rCpa1vaccine (circle), mock (triangle, GCP-MSA) and unvaccinated (square) and then they were separately challenged with approximate 90-120 viable spores isolated from one of the three *Cp* clinical isolates (C735, Silveira, or 3488) (Representative of two independent studies, n=8 mice per group). Mouse body weight at the time of challenge was set as 100%. Data points represent average of percentage body weight measured daily for 14 days. (D) Colony forming unit (CFUs) were determined by plate culture of whole organ (lung and spleen) homogenates of unvaccinated, mock and vaccinated mice at 14 DPC (n=8 per group). The data are presented as Whisker-box plots of CFUs (Log_10_) detected on plate cultures. Error bars represent the ±SEM per time point. Asterisks and daggers indicate statistically significant differences between CFU values of the GCP-rCpa1-vaccinated versus non-vaccinated mice (*, *P* < 0.05) and the vaccinated versus mock groups (†, *P* < 0.05) as determined by Ordinary one-way ANOVA comparing percent weight change at 14 DPC. Statistical differences in fungal burden were conducted using Kruskal-Wallis test.

## Methods

### Fungal culture

*Coccidioides posadasii* isolates (C735, Silveira and RMSCC 3488) and *C. immitis* isolates (RS and RMSCC 2394) were all obtained from patients with confirmed coccidioidomycosis and maintained in a BSL 3 laboratory at the University of Texas at San Antonio (UTSA) (8, 22). *Coccidioides* were grown on GYE agar plates (1% glucose, 0.5% yeast extract, and 1.5% agar) at 30 °C for 2-4 weeks to produce spores as previously reported (23).

### DNA sequence analysis

Primer 3™ software was used to design primers for amplification of the 6 rCpa1 antigens gene construct (GenBank accession AVH85517.1) as described in the **Supplementary Table 1**. Genomic DNA samples were isolated from *Coccidioides* cultures or The Fungal Testing laboratory at UT Health San Antonio. The targeted genes were amplified using a standard PCR protocol with a high-fidelity *Taq* polymerase (Life Technologies; Cat# 11304011). The PCR products were sequenced and analyzed using BioEdit™ software. The translated aa sequences of these 6 antigens were aligned using ClustalW multiple sequence alignment.

### Mice

All animal experiments were conducted following NIH guidelines and in compliance with PHS Policy on Humane Care and Use of Laboratory Animals. Protocol #MU004 approved via IACUC at UTSA. Male and female < 8-week old HLA-DR4 (DRB1*0401) transgenic mice in a C57BL/6 genetic background were bred in house (24). WT C57BL/6 mice were purchased from the NCI/CRL. Mice were euthanized humanely by an overdose isoflurane inhalation followed by cervical dislocation. Tissues and organs were collected postmortem.

### Vaccination protocol, animal challenge, and evaluation of protection

The rCpa1 construct was loaded into glucan-chitin particles (GCPs) as previously reported (18). Each dose of vaccine contained 10 μg rCpa1, 200 μg yeast tRNA and 25 μg mouse serum albumin (MSA) as a trapping matrix and 200 μg of GCPs in 200 μl PBS. Mice were subcutaneously immunized 3 times in the abdominal region at 2-week intervals followed by an intranasal challenge with 90- 150 viable spores prepared from *C. posadasii* (C735, Silveira and 3488) or *C. immitis* (RS and 2394) in 35 μl of PBS at 3 weeks after the final vaccination as described previously (19). GCP- MSA contained the same components of the vaccine except rCpa1 was used as an adjuvant control. Mice were sacrificed at 14 days post-challenge for determining fungal burden as previously described (19, 25).

### FACS analysis

Pulmonary leukocytes were isolated as previously reported (19, 25). Total cell numbers of each sample were determined using a hemocytometer. An aliquot of cells (5 ×10^5^) was labeled with a viability dye (BioLegend Zombie Aqua; Cat# 423101) and fluorochrome-conjugated antibodies for enumerating of activated T cells that expressed high levels of CD44. The antibody cocktail included FITC-CD45 (BioLegend Cat# 103108), BV421-CD44 (BioLegend Cat# 103040), BV711-TCR-β (BioLegend Cat# 1090243), APC-Fire750-CD4 (BioLegend Cat# 100460), and PeCy-7-CD8 (BioLegend Cat# 100722). IFN-γ- and IL-17A- producing T cells were determined by intracellular cytokine assays as previously described (19, 23). Briefly, cells were permeabilized and labeled with the described cocktail plus APC-IFN-γ (BioLegend Cat# 505810) and PE-IL-17A (BioLegend Cat# 506903). Data were acquired by a BD LSRII cytometer and analyzed using FlowJo software version 11.

### Statistical analyses

Fungal burden (CFUs) between two groups were analyzed via Mann-Whitney ranking test (23, 26). Ordinary one-way ANOVA was used to analyze percent weight change and Kruskal-Wallis analysis was used to analyze fungal burden and total cell numbers of cytokine-producing T cells in the lungs (19, 23). A *p* value of equal or less than 0.05 was considered statistically significant. Error bars represent the ±SEM for percent weight change and total cell number FACS analysis. Asterisks and daggers indicate statistically significant differences between GCP-rCpa1-vaccinated versus non-vaccinated mice (*) and the vaccinated versus mock groups (†).

## Results

### Variants of vaccine antigens among clinical isolates of *Coccidioides*

The multivalent rCpa1 antigen construct (GenBank: AVH85517.1; 586 aa in length) is derived from *C. posadasii* isolate C735. The rCpa1 antigen contains 3 *Coccidioides* antigens (Ag2/Pra_27-132_, Cs-Ag_138-283_ and Pmp1_288-454_) and 5 human MHC II-binding peptides derived from Pep1, Amn1 and Plb antigens (18). Sequence analysis revealed 7 aa substitutions in the rCpa1 construct among the orthologs of *C. posadasii* (*Cp*) and C. *immitis* (*Ci*) (**Table 1**). Six of the 7 aa substitutions are located on the whole length antigens (Ag2/Pra, Cs-Ag, and Pmp1). The other substitution is located on the human Plb-P6 epitope. Only 4 to 6 coccidioidal orthologs of each antigen in the rCpa1 construct were available in the GenBank. Therefore, we validated these differences by sequence analysis of the PCR amplicons using the gene-specific primers listed in (**Supplementary Table 1**) for additional 39 and 17 clinical isolates that were typed to be *Cp* or *Ci*, respectively. Our results revealed that the translated aa sequences of these antigens were identical among the 39 isolates of *Cp*. Similarly, these antigens were indistinguishable among the 17 isolates of *Ci*. Overall, these data have validated that these 7 aa substitutions between *Cp* and *Ci* (**Table 1**).

**Table 1.** Amino acid substitutions of coccidioidal vaccine antigens and peptides in the rCpa1 construct between *C. posadasii* (*Cp*) and *C. immitis* (*Ci*)

| Antigen/<br>epitope | *Cp*<sup>a</sup><br>(39 isolates)<sup>b</sup> | *Ci*<sup>a</sup><br>(17 isolates)<sup>c</sup> | GenBank no.<br>*Cp* (aa position)<sup>d</sup> | GenBank no.<br>*Ci* (aa position)<sup>d</sup> |
| --- | --- | --- | --- | --- |
| Ag2/Pra | D<sub>117</sub> | E<sub>117</sub> | EER27008.1<br>(91) | XP_001240075.1<br>(91) |
| Cs-Ag | A<sub>164</sub> | T<sub>164</sub> | AAN73410.1<br>(27, 41) | XP_001247410.1<br>(27, 41) |
|  | P<sub>178</sub> | A<sub>178</sub> |  |  |
| Pmp | M<sub>424</sub> | I<sub>424</sub> | ABB42829.1<br>(136, 142, 163) | XP_001241932.1<br>(136, 142, 163) |
|  | Q<sub>430</sub> | K<sub>430</sub> |  |  |
|  | I<sub>451</sub> | F<sub>451</sub> |  |  |
| Plb-P6 | W<sub>576</sub> | F<sub>576</sub> | ABA12208.1<br>(522) | XP_001241137.2<br>(522) |

<sup>a</sup>One letter amino acid abbreviation with a subscript number indicating position in the multivalent rCpa1 construct (GenBank accession AVH85517.1). Amino acid sequences of each antigens were aligned using Multiple Sequence Alignment (ClustalW) tool.
<sup>b</sup>Geographic distribution of *Cp* isolates: California (n=4), Arizona (n=6), New Mexico (n=1), and non-determined (n=28)
<sup>c</sup>Geographic distribution of *Ci* isolates: San Joaquin Valley, California (n=4) and Non-San Joaquin Valley (n=13).
<sup>d</sup>Corresponding positions of the aa substitution in the reference sequence of *Cp* or *Ci*.

### The GCP-rCpa1 vaccine is protective against multiple clinical isolates of *Cp*

Population genomic analysis of *Coccidioides* revealed extensive genome hybridization among clinical isolates of *Cp* and *Ci* (5, 27). Isolates C735 and Silveira pose typical *Cp* reference genomes, while RMSCC 3488 is a hybrid, with most of the *Cp* DNA sequence (personal communication with Dr. Bridget Baker). All three *Cp* isolates have identical protein sequences in the rCpa1 antigen (18). First, we evaluated the protective efficacy of the GCP-rCpa1 vaccine against a pulmonary challenge with one of the three *Cp* isolates in a susceptible C57BL/6 murine model. Mice were subcutaneously vaccinated and intranasally challenged with a lethal dose of *Coccidioides* spores as described in (**Fig. 1B**). Unvaccinated mice or mice injected with GCP adjuvant loaded with mouse serum albumin (GCP-MSA) served as negative and mock controls, respectively (18, 19). Mice were then left to rest for three weeks prior to the challenge to avoid non-specific response by innate immunity to the vaccine. Mice vaccinated with the GCP-rCpa1 vaccine and then challenged with one of the three selected *Cp* isolates maintained their body weight during the first 14 days post-challenge, while unvaccinated and mock (GCP-MSA) mice significantly reduced body weight (**Fig.1C**). The decline in the weight of unvaccinated and mock mice correlateed with a significant increase in fungal burden of the lungs and the spleens compared to their vaccinated mouse counterparts (*, *P* < 0.05) as well as mock-immunized compared to vaccinated mice (†, *P* < 0.05) (**Fig. 1D**). Notably, the GCP-MSA mock mice had less reduced bodyweight that concurred with significantly reduced fungal burden after challenge with *Cp* 3488 spores compared to unvaccinated mice (*, *P* < 0.05). These data suggest that GCP- MSA could stimulate partially protective immunity against *Cp* 3488 isolate. Therefore, the GCP- rCpa1 vaccine confers immune protection for all 3 tested *Cp* isolates.

### GCP-rCpa1 vaccine cross-protected C57BL/6 mice against *Ci* isolates

Amino acid substitutions may contribute to differences in antigenicity; therefore, we evaluated the protective efficacy of GCP-rCpa1 vaccine by C57BL/6 mice that were challenged with *Ci* isolates. RS isolate has a reference genome of *Ci*, while RMSCC 2394 is a *Cp* and *Ci* hybrid with *Ci*-type of protein sequences in the vaccine antigens. Mice were vaccinated and challenged with the same protocol in (**Fig. 1B)**. *Cp* C735 served as a reference isolate. GCP-MSA-immunized and nonvaccinated mice succumbed to coccidioidomycosis and significantly reduced body weight after a pulmonary challenge with the 2 tested *Ci* isolates despite a slower rate compared to those infected with *Cp* C735 isolate (**Fig. 2A**). Overall, mice vaccinated with GCP-rCpa1 and separately challenged with a potentially lethal dose of spores isolated from either RS or 2394 isolates significantly reduced fungal burden in the lungs and spleen compared to unvaccinated (*, *P* < 0.05) and mock GCP-MSA-injected counterparts (†, *P* < 0.05), like those challenged with C735 isolate (**Fig. 2B**). Overall, our data suggest that the GCP-rCpa1 vaccine is cross-protective against both *Cp* and *Ci* species.

**Figure 2.**
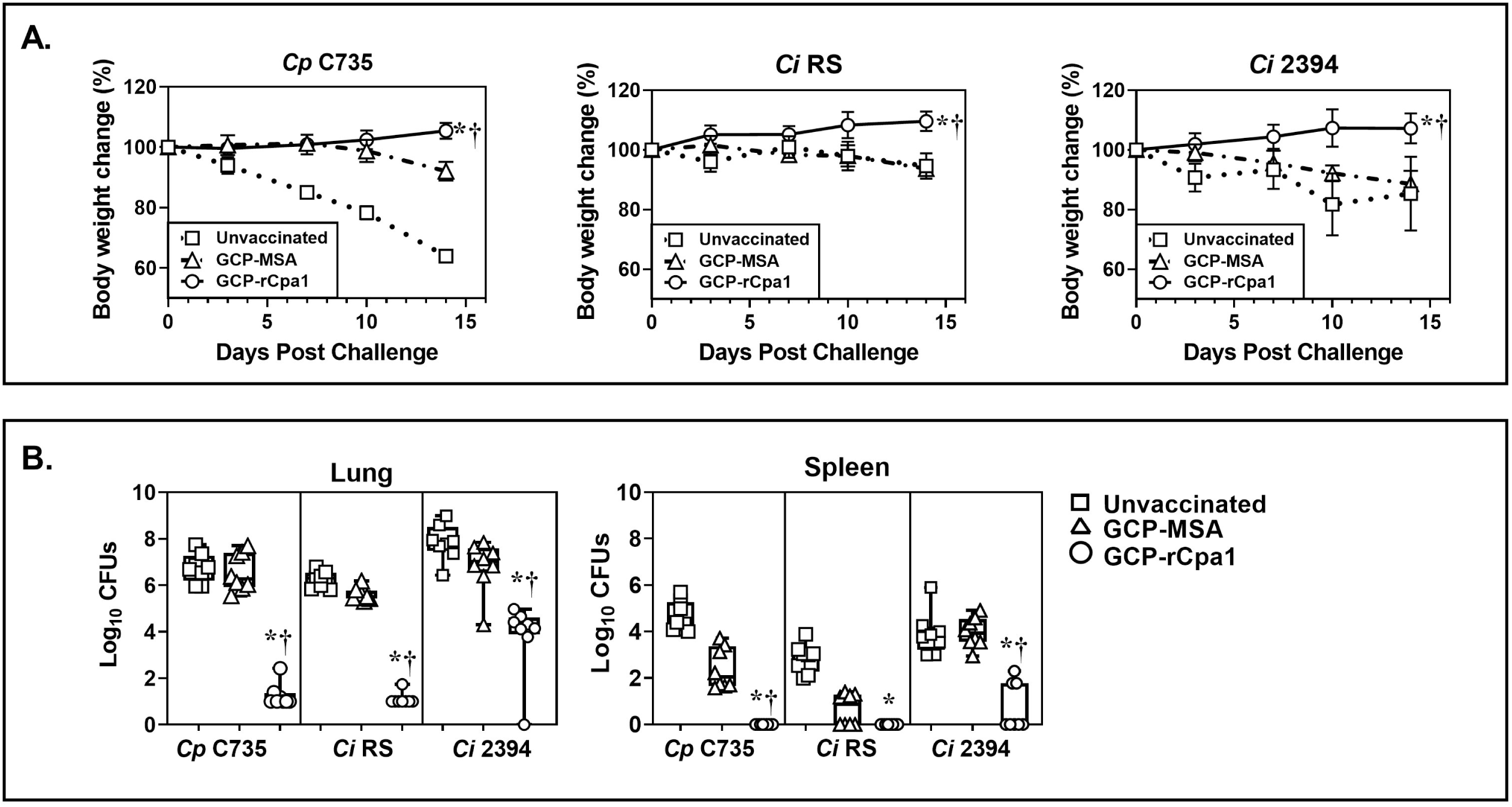
The GCP-rCpa1 vaccine confers cross-protection against isolates of *Ci*. C57BL/6 mice were vaccinated with the GCP-rCpa1 vaccine (circle), mock immunized with GCP-MSA (triangle) and unvaccinated (square) using the same schedule shown in **Figure 1B**. (A) Representative daily body weight changes (%) of two independent studies using the three groups of mice that were separately challenged with approximate 90-120 spores isolated from 2 *Ci* clinical isolates (RS and 2394). Mouse body weight at the time of challenge was set as 100% (n=8 mice per group). Data points represent average of percentage body weight measured daily for 14 days. Mice challenged with *Cp*-C735 served as a reference for comparison. (B) CFUs were determined by plate culture of lung and spleen homogenates of unvaccinated, mock and vaccinated mice at 14 DPC (n=8 mice per group). The data are presented as Whisker-box plots of CFUs (Log_10_). Unvaccinated mice, which were challenged with *Ci*-RS posed significantly reduced CFUs in the lungs and spleen compared to isolate *Cp*-C735. These data suggest that *Ci*-RS is less virulent. All groups of GCP-rCpa1-vaccinated mice had reduced CFUs in the lungs and spleen compared to the mock groups. Asterisks and daggers indicate statistically significant differences between CFU values of the GCP-rCpa1-vaccinated versus non-vaccinated mice (*, *P* < 0.05) and the vaccinated versus mock groups (†, *P* < 0.05) as determined by Ordinary one-way ANOVA comparing percent weight change at 14 DPC. Statistical differences in fungal burden were conducted using Kruskal- Wallis test.

### Vaccination with GCP-rCpa1 resulted in early induction of a mixed Th1 and Th17 response against both *Cp* and *Ci*

Studies have demonstrated that protective efficacy against coccidioidomycosis is associated with an early response via T helper cell expansion, particularly Th1 and Th17 cells (18, 19, 23, 28). We then compared pulmonary Th-cell profiles in the lungs of C57Bl/6 mice after challenge with the autologous isolate (C735) and other isolates of *Cp* and *Ci* at 7 and 14 DPC. Gating strategies for subsets of total CD4^+^ Th cells (Th1 and Th17) were based on the differential expression of CD45^+^, CD4^+^, CD8^+^, IFN-γ^+^, and IL-17A^+^ as shown in **(Supplementary Fig. 1)**. Both Th1 and Th17 cells were significantly elevated for GCP-rCpa1- vaccinated and *Cp*-C735 challenged mice compared to unvaccinated and mock counterparts at 7 and 14 DPC (**Fig. 3A**; *, *P* < 0.05 for vaccinated versus mock and †, *P* < 0.05 for vaccinated versus unvaccinated). Interestingly, GCP-rCpa1-vaccinated C57BL/6 mice only significantly elevated Th17 cells at 7 and 14 DPC after challenged with Silveira spores (**Fig. 3B**; *, *P* < 0.05). Furthermore, mice that were challenged with 3488 had significantly increased amounts of Th17 cells at 14 DPC (**Fig. 3C**).

**Figure 3.**
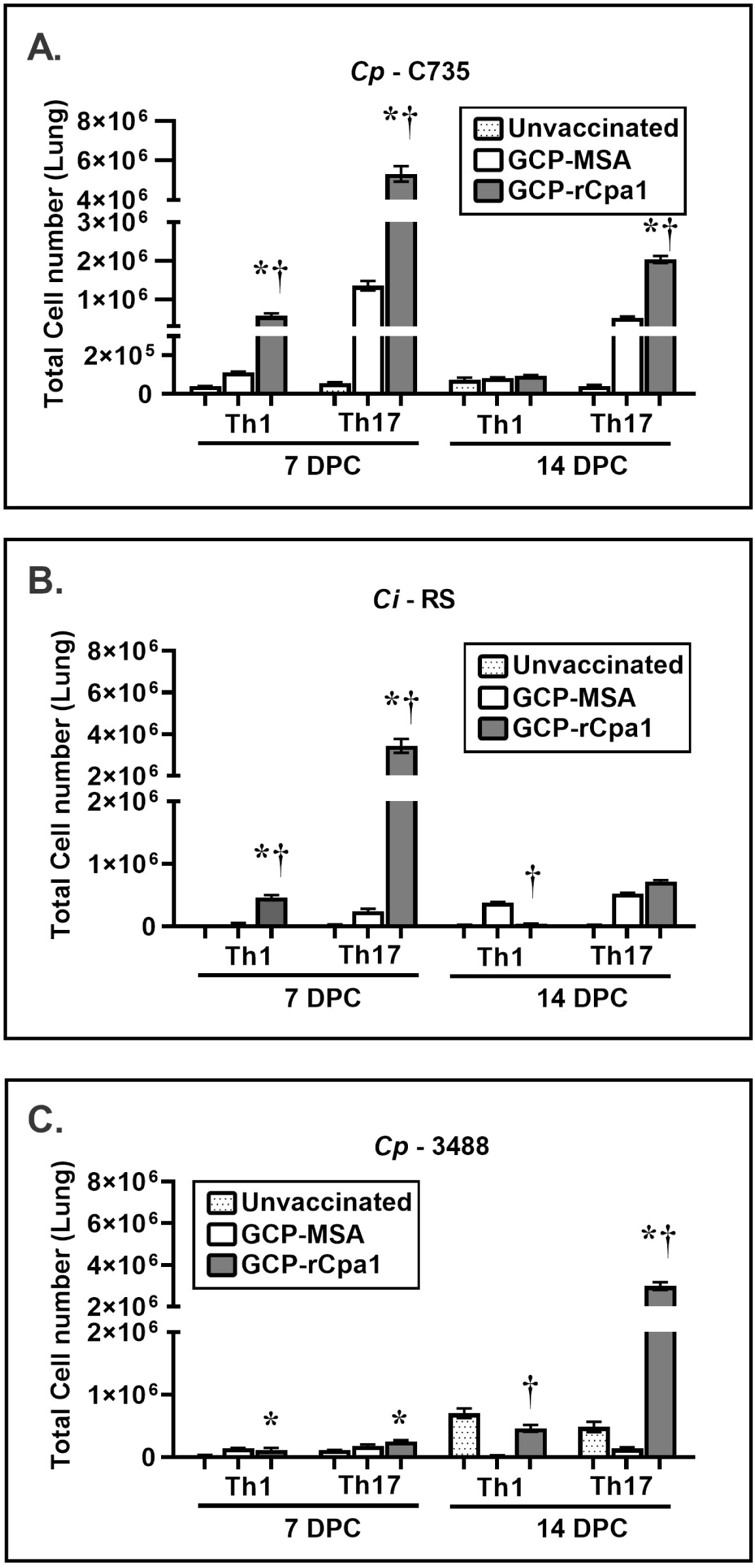
The GCP-rCpa1 vaccine induced acquisition of Th1 and/or Th17 cells in the lungs of C57BL/6 mice that were challenged with *Cp* isolates. Mice were vaccinated with GCP-rCpa1 vaccinated (black bars), injected with GCP-MSA (mock; white bars) or unvaccinated (dotted bars) using the protocol shown in **Figure 1B**. Three-weeks after the final vaccination, mice were separately challenged with a lethal dose of spores isolated from *Cp* C735 (A), *Cp* Silveira (B) and *Cp* 3488 (C). Pulmonary IFN-*γ*- and IL-17A-expressing Th1 and Th17 cells were evaluated using intracellular cytokine assays and they were gated as shown in **Fig. 3**. The absolute numbers (A to C) gated, specific cytokine producing cells per lung were determined at 7 and 14 DPC. Asterisks and daggers in panels A to C indicate significantly higher absolute numbers of the respective T-cell phenotypes in lungs of the GCP-rCpa1-vaccinated versus non-vaccinated mice (*, *P* < 0.05) and the vaccinated versus mock groups (†, *P* < 0.05). Four mice per group, per time point were used. The results are presented as mean values ±SEM and analyzed using Ordinary one-way ANOVA.

Subsequently, we profiled pulmonary CD4^+^ Th cells of C57Bl/6 mice that were challenged with *Ci* isolates. Results revealed that numbers of both Th1 and Th17 were significantly elevated in GCP-rCpa1-vaccinated mice that were separately challenged with *Ci* RS and 2394 isolates at 7 DPC (**Fig. 4A** and **4B**). These data were compared to the cytokine profiles of mice challenged with *Cp* C735 (*, *P* < 0.05) (**Fig. 3A**). Although we noted an increase of Th1 cells in the GCP- MSA control mice at 14 DPC post-challenge with *Ci* RS isolate, the increase did not correlate with fungal clearance (**Fig. 2B**). Therefore, elevated Th1 and Th17 in the lungs of the GCP- rCpa1-vaccinated mice during the first 14 DPC post-challenge are associated with dampened fungal burden, despite the kinetics of these Th responses to a pulmonary challenge with the variable isolates were not identical.

**Figure 4.**
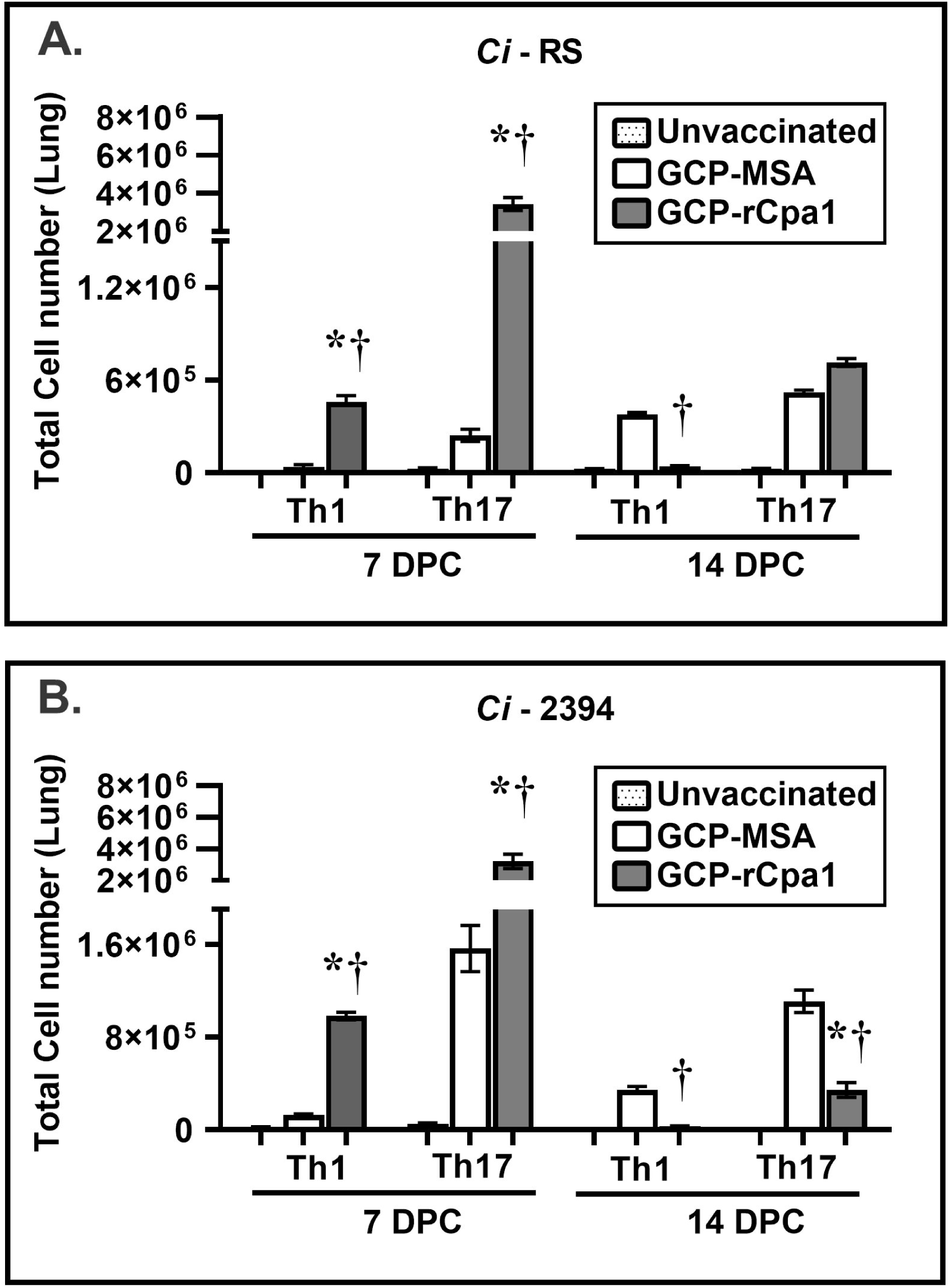
The GCP-rCpa1 vaccine induced acquisition of Th1 and/or Th17 cells in the lungs of C57BL/6 mice that were challenged with *Ci* isolates. Pulmonary Th1 and Th17 in the lungs of *Ci*-RS (A) and *Ci*-2394 were enumerated as described in Materials and Methods. Statistical significance for both Th1 and Th17 responses were observed in both *Ci* isolates (RS and 2394) at 7 DPC comparing unvaccinated to GCP-rCpa1 vaccinated mice. Total cell numbers of both Th1 and Th17 cells were significantly elevated in the GCP-rCpa1 vaccinated mice compared to mock and control mice at 7 DPC. (*, *P* < 0.05), GCP-rCpa1-vaccinated versus unvaccinated mice; (†, *P* < 0.05) GCP-rCpa1-vaccinated versus mock mice. Four mice per group, per time point were utilized. The results are presented as mean values ±SEM and analyzed using Ordinary one-way ANOVA.

### GCP-rCpa1 vaccine conferred protection for HLA-DR4 transgenic mice against both *Cp* and *Ci*

We have reported that HLA-DR4 mice are highly susceptible to a pulmonary challenge with *Cp* C735 spores. Approximately 30% of HLA-DR4 mice can approach moribund at around 11 ± 2 days post-challenge. Nevertheless, vaccination with the live attenuated (ΔT) and the GCP- rCpa1 vaccines have greatly reduced fungal burden in the lungs of HLA-DR4 mice at 9-14 DPC (29). In this study, we evaluated protective efficacy and immune responses of the GCP-rCpa1 vaccine against *Cp* 3488 and *Ci* 2394 at 9 DPC. HLA-DR4 mice challenged with *Cp* C735 served as a reference. All three isolates (*Cp*-C735, *Cp*-3488, and *Ci*-2394) are highly virulent in the C57BL/6 model of coccidioidomycosis (**Fig. 1** and **2**). HLA-DR4 mice underwent the same vaccination regimen described in (**Fig.1B)**. All three groups of vaccinated mice that were separately challenged with one of the tested isolates significantly reduced pulmonary fungal burden at 9 DPC compared to unvaccinated and mock mice, respectively (**Fig. 5A**; *, *P* < 0.05). Correspondingly to C57BL/6 mice, GCP-MSA adjuvant provided partial protection for HLA- DR4 mice against *Cp* 3488 isolate as the mock mice significantly reduced fungal burden in the lungs compared to unvaccinated mice (**Fig. 5A**). Furthermore, both Th1 and Th17 cells were significantly elevated in the vaccinated mice against *Cp*-C735 and *Cp*-3488 compared to unvaccinated and mock counterparts (**Fig. 5B** and **5C**). There was a trend of increased Th17 cells in the lungs of vaccinated mice against Ci-2394 infection compared to the mock mice, but not significant (**Fig. 5D**). These data suggest that the GCP-rCpa1 vaccine confers immune protection for HLA-DR4 mice against *Cp* and *Ci* by activating a mixed Th1 and Th17 response (†, *P* < 0.05).

**Figure 5.**
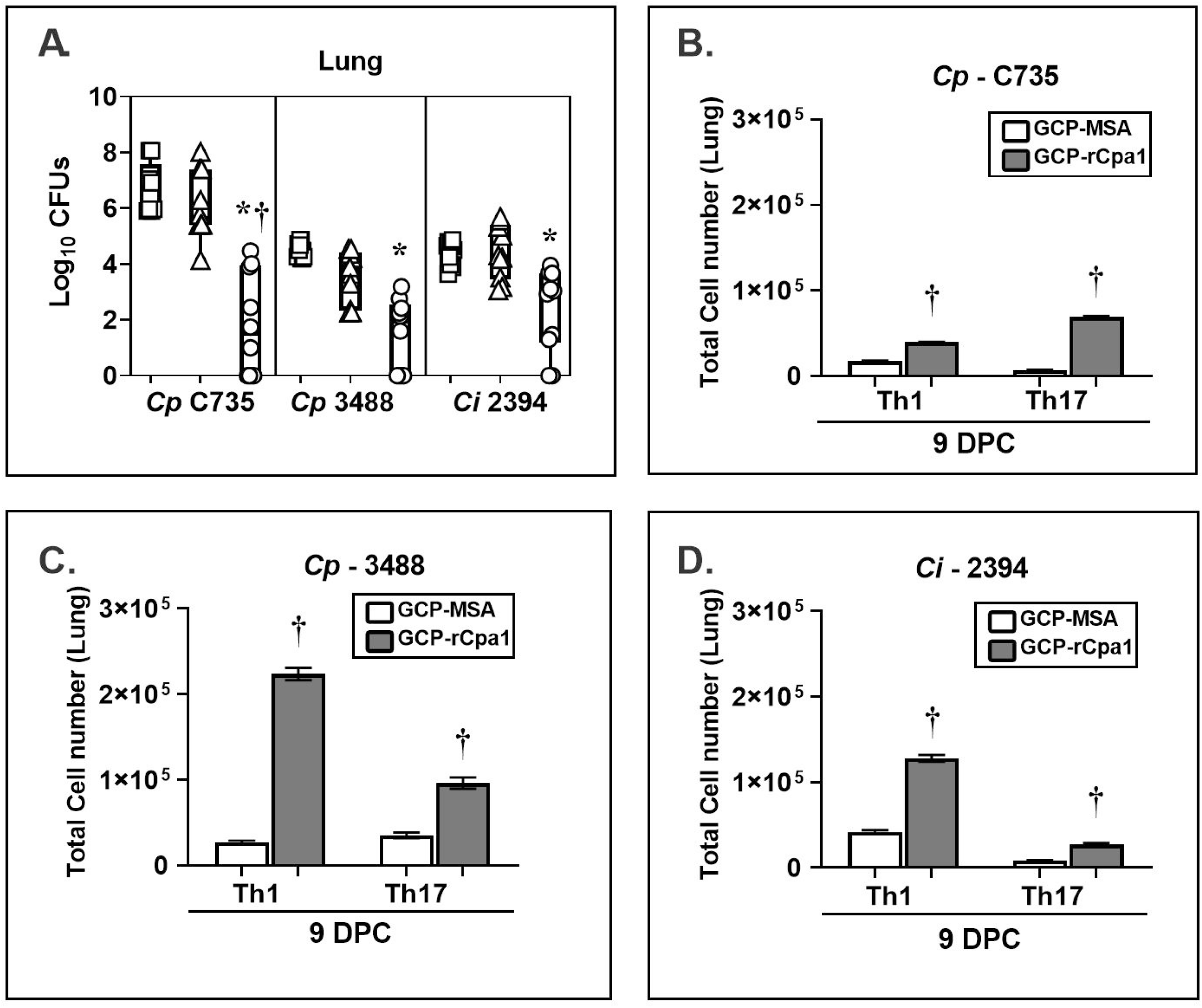
The GCP-rCpa1 vaccine elicits cross protection and a mixed Th1 and Th17 response in the lungs of HLA-DR4 transgenic mice. HLA-DR4 (DRB1*0401 allele) transgenic mice were immunized and then separately challenged with *Cp* C735, *Cp* 3488 and *Ci* 2394. HLA-DR4 mice were subcutaneously vaccinated with the GCP-rCpa1vaccine (circle), mock (triangle, GCP-MSA) and unvaccinated (square) and then they were separately challenged with approximate 90-120 viable spores isolated from *Cp* isolates (C735, or 3488) or *Ci* (2394) (10 mice per group). Whisker-box plots presented CFUs (Log_10_) values from whole lungs at 9 days post challenge (A). Pulmonary Th1 and Th17 cells were evaluated by intracellular cytokine staining assays. Asterisks and daggers indicate statistically significant differences between CFU values of the GCP-rCpa1-vaccinated versus non-vaccinated mice (*, *P* < 0.05) and the vaccinated versus mock groups (†, *P* < 0.05) as determined differences in fungal burden was conducted using Kruskal-Wallis test. (B) Both Th1 and Th17 cells were significantly elevated for GCP-rCpa1-vaccinated mice compared to mock mice (GCP-MSA) that were challenged with *Cp* C735 (B) and *Cp* 3488, (C) and *Ci* 2394 (D). Ordinary one-way ANOVA was used to evaluate vaccinated versus mock groups (†, *P* < 0.05), 8 mice per group were used for FACS studies.

## Discussion

We demonstrated that GCP-rCpa1 offers protection for C57BL/6 and HLA-DR4 transgenic mice against both *C. posadasii* and *C. immitis*, despite the 7 amino acid substitutions present in the proteins and peptides of the multivalent rCpa1 antigen between the two fungal species. The principal advantage of multivalent subunit vaccines is its safety, its components only contain microbe-specific proteins or synthetic peptides, a chemically-defined adjuvant, and a delivery system. Recombinant vaccine antigens can be engineered to overcome antigenic variability and remove human homologous sequences using sequence-based prediction tools and genomics/bioinformatics analysis. Immuno-bioinformatics analysis using ProPred algorithm (http://crdd.osdd.net/raghava/propred/) contains information for 51 alleles of human leukocyte antigen type HLA-DR and a privileged MHC epitope prediction software for commonly used laboratory mice and 61 human HLA alleles (IoGenetics LLC, Dr. Jane Homan personal communication) was applied to predict potential MHC II-binding epitopes in the rCpa1 antigen. Results suggest that six of the seven substitutions are not located on the predicated MHC II binding epitope regions of humans nor C57BL/6 mice. The substitution (W_576_ > F_576_) in the Plb- P6 epitope may not alter the immunogenic property and protective efficacy of the rCpa1 antigen for C57BL/6 mice since it is not recognized by their MHC II molecules (I-Ab allele) (18). Additionally, this substitution does not alter the protective efficacy of the rCpa1 antigen for both C57BL/6 and HLA-DR4 mice against either *C. posadasii* or *C. immitis* infection as shown in (**Fig. 1, 2**, and **5)**.

Successful vaccines against *Coccidioides* infection should confer protection against a broad spectrum of clinical isolates (14). Both species are estimated to diverge around 5 million years ago (5, 8), sharing major virulence determinants and structural genes, although comparison of their genomes revealed some variation in aa sequences of their translated proteins (1, 27). *Cp* mycelia grows significantly faster at 37 ºC when compared to *Ci*, despite growth rates of parasitic spherules for those isolates at human physiological temperature have not yet been explored (30). Clinical isolates of *Cp* and *Ci* are all highly virulent in experimental murine models of coccidioidomycosis that LD_100_ doses for most isolates are lower than 50 viable spores (4, 31). We have evaluated protective efficacies of the GCP-rCpa1 vaccine against 3 isolates of *Cp* (C735, Silveira, and RMSCC 3488) and 2 isolates of *Ci* (RS and RMSCC 2394). *Cp*-C735 was isolated from a patient who visited the VA hospital in the 1970s in San Antonio, Texas. *Cp*- Silveira was isolated from the first reported case of Valley Fever from a patient who lived in Arizona in 1894 (32). *Cp*-RMSCC 3488 isolate was acquired from a patient who lived in southern/central Mexico. Isolates RS and RMSCC 2394 of *Ci* were obtained from patients residing in central California and the southern California/northern Mexico regions, respectively. Overall, these 5 isolates present a wide geographic distribution from Arizona, California, Texas, and central Mexico (33), and were obtained at various time point during the course of history of the disease (4) (27). RMSCC 3488 and RMSCC 2394 genomes pose hybrid DNA sequences from two reference genome databases of *Cp* C735 and *Ci* RS, while they are typed to be *Cp* and *Ci*, respectively(7). Concurringly, DNA sequence analysis of the orthologous proteins included in the rCpa1 antigen demonstrated that RMSCC 3488 has *Cp*-type of the vaccine antigen and RMSCC 2394 has *Ci*-type. Remarkably, we found vaccination with GCP-rCpa1 provided almost identical levels of protection for C57/BL6 mice against these 5 isolates of *Coccidioides*. In each case, the pulmonary fungal burden was significantly reduced by four Log10 orders at 14 DPC.

Furthermore, vaccination with GCP-rCpa1 also had significant reduction of fungal burden for highly susceptible HLA-DR4 mice against *Cp* and *Ci* (2-4 Log_10_) at 9 DPC. Taken together, these results support our hypothesis that the GCP-rCpa1 has cross-protection activity against various isolates of *Cp* and *Ci*. These data also suggest that the 7 aa substitutions in the rCpa1 vaccine antigen do not impact protective efficacy against both species of *Coccidioides*.

The strongest immune correlative of protection against pulmonary coccidioidomycosis is in the early acquisition of Th1 and Th17 cells that produce IFN-g and IL-17 in the lungs, respectively (23, 25, 34). Both Th1 and Th17 cells play a central role in defense against *Coccidioides* infection (18, 19, 31, 35, 36). The genetically resistant DBA/2 mice developed a Th1-biased response, with an early induction of IFN-g, whereas susceptible BABL/c mice showed an early secretion of the Th2 cytokine, IL-4 (35). Previous studies evaluating the immunogenicity of GCP-rCpa1 vaccine via cytokine recall assay demonstrated that following at least a 2-week resting period, mock immunized group (GCP-MSA) were unable to elicit non-specific T-helper response. While GCP-rCpa1 immunized mice elicited a strong mixed Th1/Th17 response with little Th2 response by IL-4 production (18). Neutralization of IL-12, a critical cytokine for differentiation of Th1 cells, in BDA/2 mice led to increased susceptibility to *Cp* Silveira infections (36). Furthermore, C57Bl/6 and BALB/c mice are able to produce IL-10 at greater levels compared to DBA/2 mice. Many studies have shown IFN-g production as a correlate of vaccine-induced protection for mice against *Coccidioides* infection (4, 37-40). An attenuated live vaccine (ΔCps1) created from *Cp* Silveira isolate elicits a Th1-biased response that confers protection against a pulmonary challenge with the autologous isolate or *Ci* RS isolate (31, 41). In humans, polymorphisms in IL-12/IFN-γ signaling pathway resulted in a STAT1 gain-of-function mutations that associates with increased disease severity to *Coccidioides* infection (42). A homozygous C_186_Y mutation in the IL-12β1 receptor is associated with increased risk of disseminated coccidioidomycosis (43). Furthermore, a case report shows supplementation of antifungal agents with IFN-γ slowed disease progression, and the addition of IL-4 and IL-13 blockade with dupilumab resulted in rapid resolution of the patient’s clinical symptoms (44). Those observations support the hypothesis that Th1 response plays an important role in defense against *Coccidioides* infection.

We have previously reported that subcutaneous vaccination of IFN-γ- and IL-4-receptor knockout mice with an attenuated, live vaccine (ΔT) derived from *Cp* C735 isolate elicited protective immunity against pulmonary challenge with an autologous virulent isolate (44). In contrast, fungal burden, clearance, and survival are significantly compromised in mice defective in IL-17A or IL-17R (23). Similarly, we demonstrated that depletion of IL-17A in mice reduces the protective efficacy of the GCP-rCpa1 vaccine against a pulmonary challenge with Cp C735 isolate (18, 19). A recent patient with disseminated coccidioidomycosis was found to have a STAT3 mutation. STAT3 mediates IL-23 signaling, which is critical for IFN-γ, IL-12, and IL-17 production (45).These observations suggest that Th17 cells and IL-17 are indispensable for vaccine immunity against *Coccidioides* infection.

Both Th1 and Th17 cells are significantly elevated against *Cp* C735 infection at 7 DPC as previously reported (**Fig. 3A**; (18, 19)). Similarly, Th1 and Th17 cells are also increased in response to a pulmonary challenge with isolates *Ci* RS and *Ci* 2394 at 7 DPC (**Fig. 4**). Notably, only Th17 cells are significantly increased in the vaccinated mice against isolates *Cp* Silveira and *Cp* 3488 at 7 and 14 DPC, respectively **(Fig. 3B** and **3C**). The GCP-rCpa1-vaccinated mice induce activation of pulmonary Th1 and Th17 cells at various levels and temporal patterns in response to a pulmonary challenge with various isolates of *Cp* and *Ci*. These differences are not associated with a respiratory challenge with *Cp* versus *Ci*, despite evaluation of only 5 isolates limits the capacity for interspecies and interspecies comparisons. The most common correlate of GCP-rCpa1 vaccine-induced protection against all 5 isolates is significantly increased acquisition of Th17 cells in the lungs. The vaccine-induced response may be dependent on the fungal isolates, challenge dose, and host variation in an immune response. Relative contribution of Th1 and Th17 cells for mice against different *Coccidioides* isolates requires further investigation.

In summary, our study supports the concept that a combination of genetic fusion techniques to create a multivalent antigen is a perspective approach for the generation of a broadly effective vaccine candidate that protects against both *Cp* and *Ci*. It is imperative to investigate multivalent vaccine formulations with broad-spectrum protection and sufficient safety in the development of a human vaccine against an orphan disease, such as coccidioidomycosis, that is caused by multiple species of fungi.

## Author Contributions

Conceptualization, AC, KDP, and CYH.; methodology, AC, KDP, YRL, HZ, GO CYH.; software, AC, KDP, YRL, HZ, CYH.; Validation, KDP, AC, YRL.; formal analysis, KDP, HZ, NW, CYH; investigation, NW, GO, CYH.; Resources, NW, GO, CYH. Original draft preparation, AC, KDP, CYH.; Writing, AC, KDP, CYH.; review and editing, AC, KDP, YRL, HZ, CYH.; visualization, CYH.; supervision, CYH.; funding acquisition, CYH. All authors have read and agreed to the published version of the manuscript.

## Acknowledgments

We thank Dr. Sandra Cardona, Manager of the Cell Analysis Core at UTSA for excellent technical assistance in flow cytometry analysis and Dr. Jieh Juen Yu at UTSA for helpful discussion.

---

**Supplementary Table 1.**
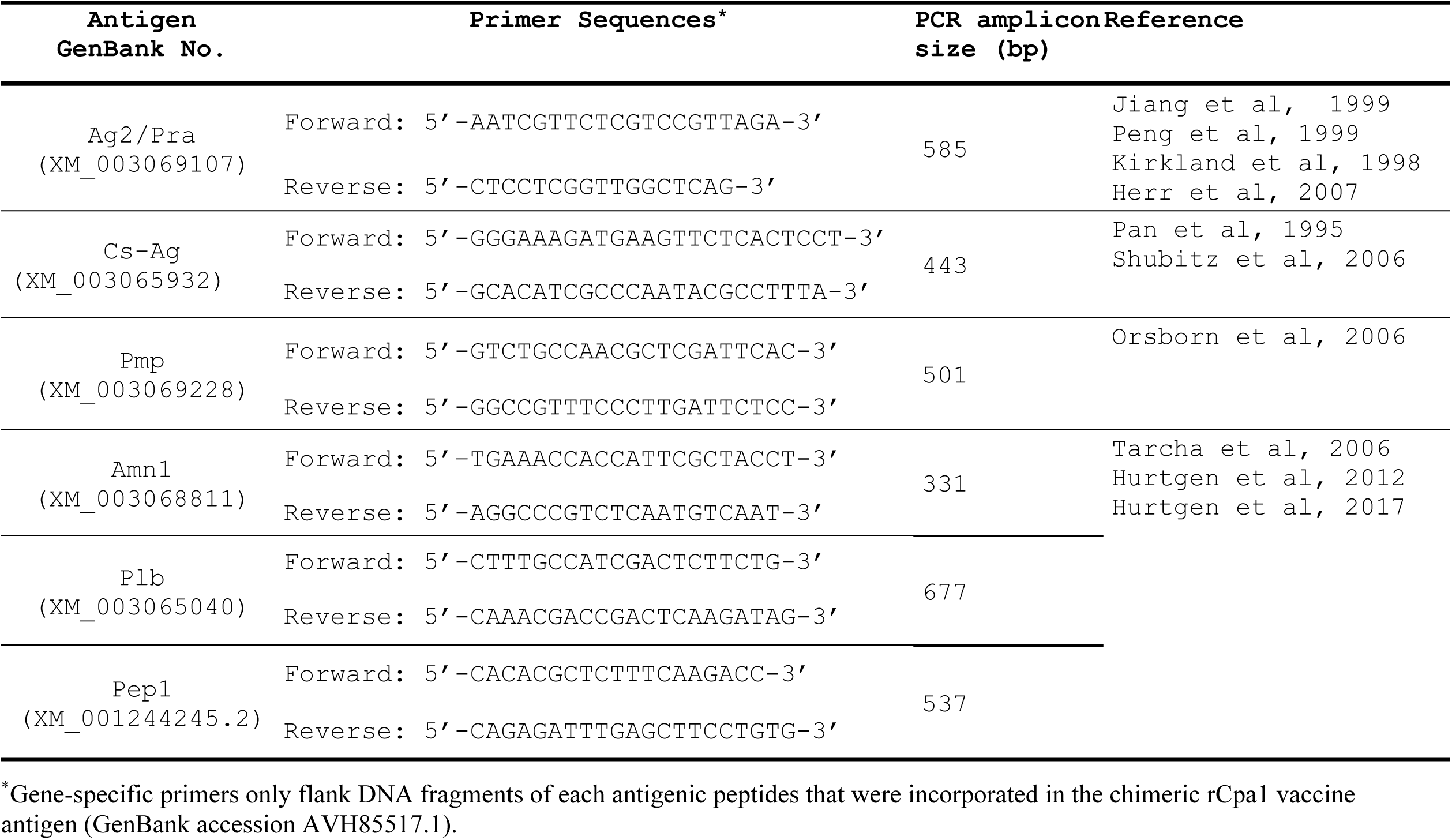
Primer sequences and predicted sizes of PCR amplicons of the *Coccidioides* antigens

